# Eco-evolutionary dynamics of clonal multicellular life cycles

**DOI:** 10.1101/2022.03.14.484302

**Authors:** Vanessa Ress, Arne Traulsen, Yuriy Pichugin

## Abstract

The evolution of multicellular life cycles is a central process in the course of the emergence of multicellularity. The simplest multicellular life cycle is comprised from the growth of the propagule into a colony and its fragmentation to give rise to new propagules. The majority of theoretical models assume selection among life cycles to be driven by internal properties of multicellular groups resulting in growth competition. At the same time, the influence of interactions between groups on the evolution of life cycles is rarely even considered. Here, we present a model of colonial life cycles evolution taking into account group interactions. Our work shows that the outcome of evolution could be coexistence between multiple life cycles or that the outcome may depend on the initial state of the population – scenarios impossible without group interactions. At the same time, we found that some results of these simpler models remain relevant: Evolutionary stable strategies in our model are restricted to binary fragmentation – the same class of life cycles which contains all evolutionarily optimal life cycles in the model without interactions. Our results demonstrate that while models neglecting interactions can capture short-term dynamics, they fall short in predicting the population-scale picture of the evolution.

## 1 Introduction

Multicellular organisms are found everywhere. In all major branches of complex multi-cellularity (animals, plants, fungi, red, and brown algae) organisms are formed by cells staying together after cell division - unlike to unicellular species, in which cells part their ways before the next division occurs. However, all organisms have to reproduce as otherwise their population cannot grow. Hence, at least occasionally, some cells must depart from a multicellular organism in order to develop into an offspring one. The combination of organism growth and reproduction constitute a clonal life cycle. Emergence of clonal multicellular life cycles was the central innovation in the earlier stages of the evolution of multicellularity. There, traits, which do not even exist for unicellular species, become crucial for long-term success of even the most primitive colony of cells. These include the number of cells in the colony, how often cells depart to give rise to new colonies, how large the released propagules are, and how many of them are produced. As the reproduction and, consequently, fitness of simple cell colonies are dependent on these traits, they immediately become subjected to natural selection, favouring some life cycles over others. Since complex multicellular life descends from those loose cell colonies, the understanding of the prior evolution of primitive life cycles is essential to our understanding of the later evolution of complex traits.

Some models of the evolution of simple multicellular life cycles assume that natural selection operates by means of growth competition. Colonies are born small but due to cell divisions they increase in size and, eventually, fragment, so the number of colonies in the population grows. The life cycle maximizing the population growth rate has a selective advantage, as it outgrows all competitors [Roze et al., 2001, Libby et al., 2014, Pichugin et al., 2017, 2019, Staps et al., 2019, Gao et al., 2019, Pichugin and Traulsen, 2020, Gao et al., 2021, Pichugin and Traulsen, 2022]. It was shown that for groups made of identical cells, some life cycles are forbidden: they cannot be the winner of growth competition under any conditions. For instance, if the fragmentation event is instantaneous and its execution does not cost anything to the group, any life cycle in which a group fragments into more than two offspring groups is forbidden. Hence, only fragmentation into two pieces can evolve under these conditions [Pichugin et al., 2017, Pichugin and Traulsen, 2020]. And indeed, the division into two pieces is, by a large margin, the most common reproductive strategy among microscopic life forms.

However, these models, due their conceptual simplicity, assume unconstrained (exponential) growth of the population, which cannot be sustained for a prolonged period of time, because both resource and space are limited. Other models consider density dependent growth [Rossetti et al., 2011, Tarnita et al., 2013, van Gestel and Nowak, 2016, Henriques et al., 2021], where the population growth decreases with the number of groups. A similar approach is the Moran birth-death process on the group level, where whenever a new group emerges, one other group dies [Traulsen and Nowak, 2006, Rodrigues et al., 2012, Simon et al., 2013, Luo, 2014, Cooney, 2019, 2020]. While the population dynamics of density-dependent population growth is vastly different from the exponential explosion found in models of unconstrained growth, these two approaches lead to identical results for life cycle evolution: As shown in Appendix 1 of [Pichugin and Traulsen, 2020], the dynamics of the fraction of a given life cycle are identical in models with unconstrained and density-dependent growth. Therefore, even in models with density-dependent growth, the evolutionary success of the life cycle is still fully determined by the population growth rates.

Nevertheless, density-dependent growth is also a simplification, as different groups may differ in their competitiveness. For instance, large cell colonies are able to block single cells from access to vital resources [Rainey and Travisano, 1998, Rainey and Rainey, 2003, Hammerschmidt et al., 2014], which may even lead to a complete extinction of solitary cells. Thus, the population dynamics of multicellular life cycles is not necessarily density dependent, but could be frequency dependent – the impact of resource limitation on the population growth depends on the composition of the population itself. Hence, the evolution of multicellular life cycles cannot always be reduced to growth competition, but may arise from eco-evolutionary dynamics.

In broader empirical perspective, frequency-dependent dynamics is found to be common among microbial populations [Levin et al., 1988, Ribeck and Lenski, 2015, Healey et al., 2016, Friedman et al., 2017]. From the perspective of the theoretical ecology, frequency-dependent evolutionary dynamics arising from interactions between diverse population members has also been considered in detail [May, 1972, Wangersky, 1978, Bomze, 1983, Huang et al., 2015, 2017, Bunin, 2017, Barbier et al., 2018, Kotil and Vetsigian, 2018, Tarnita, 2018, Farahpour et al., 2018, Park et al., 2019, 2020]. However, both empirical and theoretical ecology approaches tend to overlook frequency-dependence in the context of life cycles, where the role of an organism in interactions changes with the stage of life cycle (but see an example in [Tverskoi and Gavrilets, 2022] modelling evolution of germ-soma differentiation).

In this work, we develop a model of clonal life cycles evolution, in which cell division and fragmentation are accompanied by group competition in the form of frequency-dependent colony death rates. Groups of different sizes have different efficiency in enduring such a competition and/or applying pressure on their counterparts. Such a dependence of a group’s fate on the presence of other groups adds an ecological dimension to the question of multicellular life cycle evolution.

With this model, we study what kind of life cycles survive in the course of eco-evolutionary dynamics. We found that interactions between groups allow for situations with bi-stability or coexistence of multiple life cycles - scenarios impossible in the unconstrained growth model. When an eco-evolutionary dynamics leads to a coexistence between life cycles, a life cycle which was not able to win growth competition in the unconstrained growth model can be a member of the mixed population. Also, we found that despite the fundamental differences to the model with unconstrained growth, some of its results have a direct analogy in a much more general eco-evolutionary context. For instance, life cycles forbidden in the unconstrained model cannot be evolutionary stable strategies in the eco-evolutionary model. We also show that for a broad class of competition matrices, in which the death probability of a group only depends on the size of the opponent, the results of evolutionary competition between life cycles are exactly the same as in the model with unconstrained growth.

## 2 Model

### 2.1 Population dynamics of a single life cycle

We consider a population consisting of cell groups that grow in size and fragment, giving rise to new groups. Cells within a group of size *i* divide at rate *b*_*i*_, thus a group of size *i* grows at rate *ib*_*i*_. Groups also die due to both external environmental factors and within-population competition for resources or space. Due to the external factors, the death rate of groups of size *i* is *d*_*i*_. Frequency-dependent competition is modelled as the death of groups of size *i* upon encounter with groups of size *j* at rate *K*_*i,j*_, see Fig. 1A.

**Figure 1:**
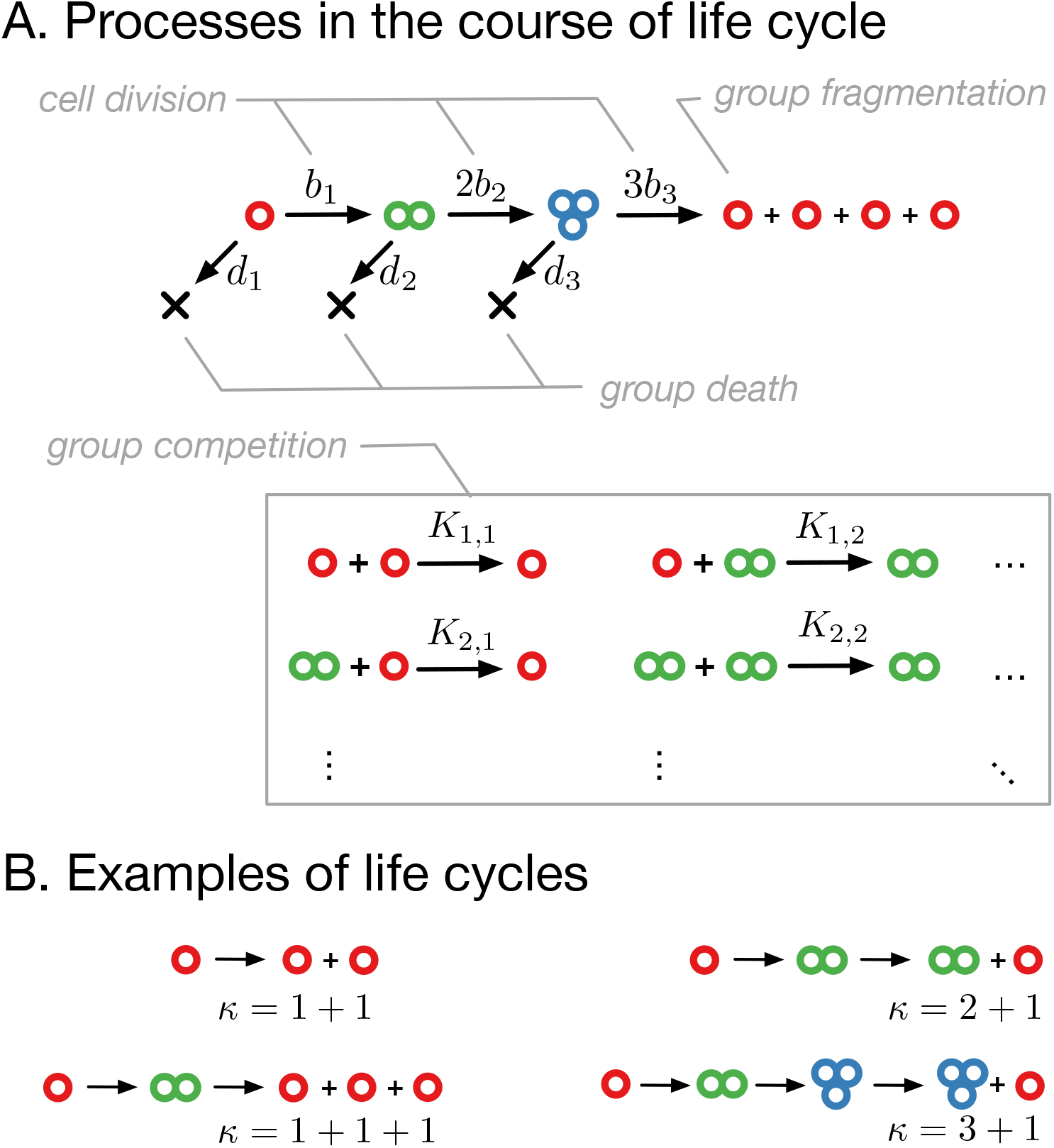
Model of clonal life cycles. **A**. There are four processes occurring in the model: groups grow by cell division, which occurs with rates *b*_*i*_, groups spontaneously die with rates *d*_*i*_, groups fragment immediately upon exceeding maturity size *m* (3 in this example), according to predefined fragmentation pattern *κ* (1+1+1+1 here), and groups of size *i* die due to the competition with groups of size *j* at rates *K*_*ij*_. **B**. Together, group growth and fragmentation constitute a life cycle, in which initially small groups grow in size and eventually fragment, giving rise to more small groups.

Whenever a group of maturity size *m* grows to *m* + 1 cells, it immediately fragments. The fragmentation always occurs by the same pattern and determines the life cycle of a population. We represent a fragmentation pattern by *κ* - a partition of a number *m* + 1. For example, the fragmentation pattern of the unicellular life cycle, in which two daughter cells always go apart, is *κ* = 1 + 1, see Fig. 1B. Other fragmentation patterns correspond to multicellular life cycles. The simplest of them are the life cycles in which groups grow up to two cells, but fragment upon reaching size three. Such a fragmentation can be performed in two ways: either detachment of a single cell, leading to the fragmentation pattern *κ* = 2 + 1, or fission into three solitary cells, *κ* = 1 + 1 + 1, see Fig. 1B. For simplicity, we assume that the cell number does not change during fragmentation (no cell loss), the sum of a fragmentation pattern *κ* is equal to *m* + 1.

If we denote the abundance of cell groups containing *i* cells as *x*_*i*_, then the dynamics of population is described by a system of differential equations

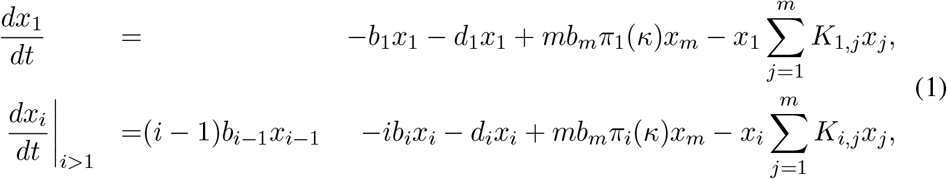

where the first two terms (*i* − 1)*b*_*i−*1_*x*_*i−*1_ − *ib*_*i*_*x*_*i*_ describe the growth of groups - the positive term represents growth from size *i* − 1 to *i* and the negative term represents growth from *i* to *i* + 1. The third term −*d*_*i*_*x*_*i*_ is the environmentally caused death. The term *mb*_*m*_*π*_*i*_(*κ*)*x*_*m*_ describes the birth of new groups of size *i* via fragmentation of larger groups, where *π*_*i*_(*κ*) is the number of groups of size *i* produced in the result of that fragmentation. Finally, 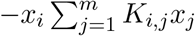 is the death of groups due to the competition between groups.

Summarising the dynamics into matrix notation, the system (1) can be written as

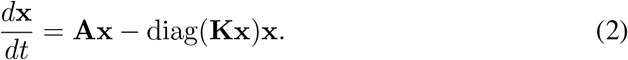

Here, **x** is the column-vector of group abundances

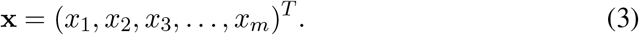

The projection matrix **A** of size *m* × *m* is given by

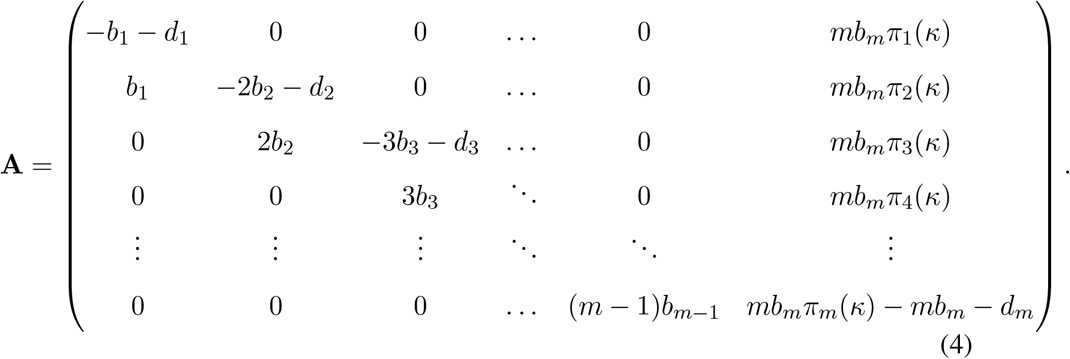

The elements of the projection matrix **A**_*i,j*_ represent changes to number of groups of size *i* due to processes occurring with groups of size *j* (but not due to interactions). Hence, the projection matrix has non-zero elements only on the main diagonal (group death and growth of groups to larger sizes), lower sub-diagonal (growth of smaller groups), and rightmost column (fragmentation at the end of the life cycle). The elements of the competition matrix **K** are given by by *K*_*i,j*_ for *i, j* = 1, …, *m*. The operation diag(·) takes an input vector of length *m* and transforms it into a diagonal matrix of size *m* × *m* with the entries of the input vector on the main diagonal.

### 2.2 Population dynamics of multiple life cycles

To investigate the eco-evolutionary dynamics of clonal life cycles, we consider a composition of several (*r*) sub-populations executing various life cycles: *κ*^(1)^, *κ*^(2)^, …, *κ*^(*r*)^. In this composite population, the cell growth (*b*_*i*_), environmentally caused (constant) group death (*d*_*i*_), and group fragmentation (*π*_*i*_(*κ*)) occur independently in each sub-population. However, frequency dependent death due to competition entangles the dynamics of the sub-populations, as groups with different life cycles growing together have to compete with each other. If we denote with 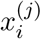 the number of groups containing *i* cells in a sub-population executing the life cycle *κ*^(*j*)^, the dynamics of the composite population is described by the differential equations:

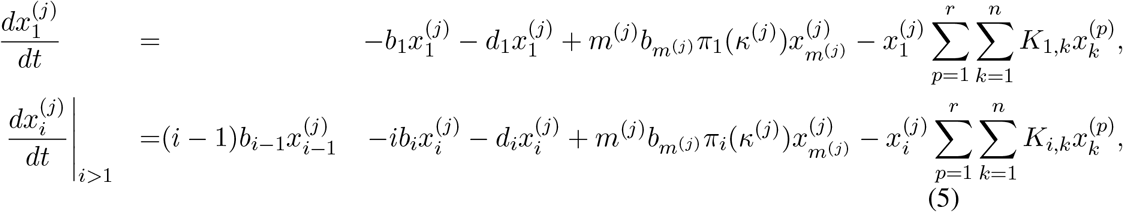

where *m*^(*j*)^ is the maturity size of the life cycle *κ*^(*j*)^, and *n* = max(*m*^(1)^, *m*^(2)^, …, *m*^(*r*)^) is the maximal maturity size in the composite population. The difference between the one life cycle system (1) and the system of multiple life cycles (5) is in the last term, where groups from every competing sub-population contribute to the frequency dependent death.

In vector form, the state of the composite population is described by a concatenation of vector states of each sub-population:

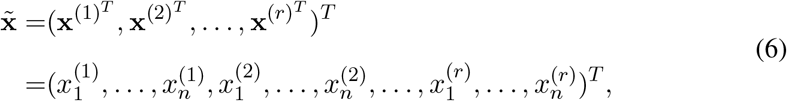

where 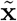 is the column-vector describing the state of the composite population, **x**^(*j*)^ is the column-vector describing *j*-th sub-population in a form (3). Note that the last entries of any 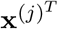 will be zero if *m*^(*j*)^ < *n*. The dynamics of the composite population in Eq. (5) can be represented in the vectorized form, similar to Eq. (2):

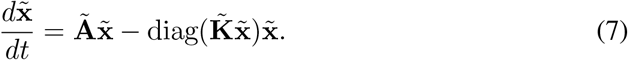

Here the composite projection matrix representing the cell growth, environmentally caused (constant) group death, and group fragmentation is a diagonal block matrix

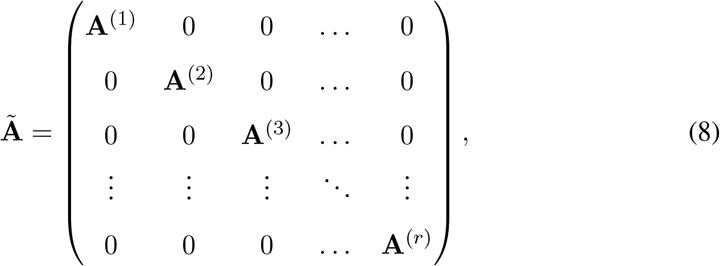

where **A**^(*i*)^ is the projection matrix of the life cycle *κ*^(*i*)^ extended to size *n* × *n* (*n* is the maximal maturity size across all competing life cycles). If the maturity size of the life cycle *i* is *m*^(*i*)^ = *n* this matrix has a form exactly as in Eq. (4). If the maturity size is smaller, *m*^(*i*)^ *< n*, then the top left *m*^(*i*)^ × *m*^(*i*)^ has the form (4), while the remaining elements are non-zero at the main diagonal and the lower subdiagonal, as dictated by Eq. (5).

The composite competition matrix 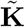 is constructed as

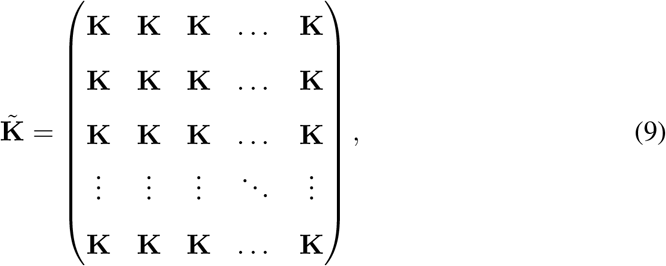

where each block **K** is a competition matrix.

### 2.3 Invasions from rare

In the general case, the investigation of the composite population dynamics given by Eq. (7) is a very complex problem without a general solution. Hence, in our study we consider a specific class of initial conditions - invasion from rare, where the composite population contains only two sub-populations: the abundant “resident” executing life cycle *κ*^(*R*)^ and rare “invader” executing different life cycle *κ*^(*I*)^. This scenario represents an emergence of a mutant in previously stable population of the resident. The population changes if this mutant is capable to invade the resident – otherwise the mutant goes extinct and the resident population remains the same.

In this scenario, the composite dynamics in Eq. (7) can be disentangled into resident and invader components. Since the invader population is small, its contribution to the frequency-dependent competition is negligible. The members of the resident population compete predominantly between themselves, so the resident dynamics is effectively a single-species scenario,

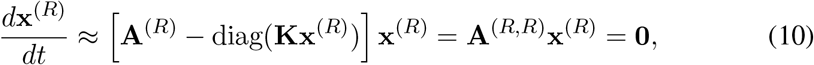

where the vector **x**^(*R*)^ represents the composition of the resident population, **A**^(*R*)^ is the projection matrix of the resident, and we introduced the self-invasion projection matrix **A**^(*R,R*)^ = **A**^(*R*)^ − diag(**Kx**^(*R*)^). Since the resident is assumed to be at a stable equilibrium in the absence of invaders, the self-invasion matrix **A**^(*R,R*)^ has an eigenvalue which is zero, and the equilibrium population composition **x**^(*R*)^ is given by the corresponding eigenvector. The resident population dynamics can be obtained by solving the non-linear Eq. (10), which in the general case can be performed only numerically.

The rare invader population also competes primarily with the resident and self-competition is negligible. Thus, its dynamics is given by

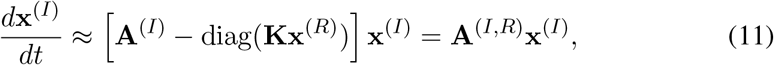

where vector **x**^(*I*)^ represents the composition of the invader population and we introduced the invasion matrix

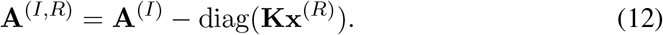

Unlike the resident dynamics, the dynamics of the invader population is linear – the invasion projection matrix **A**^(*I,R*)^ is independent from the invader population state **x**^(*I*)^. The linear dynamics of clonal life cycles has been extensively studied in previous work [Pichugin et al., 2017]. If the largest eigenvalue of the invasion matrix **A**^(*I,R*)^ is positive, then the invader population will increase in numbers, independently on its initial demography. Otherwise, the invader population goes extinct.

The assumption of a negligible impact of the invader population on competition limits the analysis to the early stages of invasion, when the invader population is small. Nevertheless, this makes it possible to investigate the stability of resident life cycles with respect to invasions.

## 3 Results

We begin with a description of the population dynamics of a single life cycle in isolation. Then, we consider the evolution of a population containing multiple life cycles in special cases of the competition matrix. Then, we consider the case of arbitrary interactions, but limit our analysis to the scenario of invasions from rare.

### 3.1 Dynamics of a single life cycle

For the simplest unicellular life cycle (*κ* = 1 + 1), where population is composed only of solitary cells, the dynamics of our model given by Eq. (2) reduces to logistic growth,

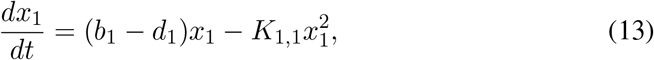

where the growth rate is equal to *b*_1_ − *d*_1_ and the carrying capacity is (*b*_1_ − *d*_1_)*/K*_1,1_.

The population dynamics of more complex life cycles also bears similarity to logistic growth. If a population is small, the competition term is small, so the population grows exponentially. While population size increases, so does the competition term – group mortality rises and the overall population growth slows down. The growth stops when the population reaches a stationary state **x**\*, where

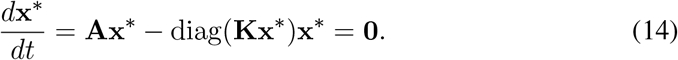

Numerical simulations show that an isolated life cycle always comes to the same stationary state **x**\* from any initial distribution of group sizes.

The equilibrium population composition **x**\* is shaped by the balance between group growth (*b*_*i*_), death (*d*_*i*_), and competition (*K*_*ij*_). For an arbitrary case, we cannot present an analytical solution for **x**\*. However, in the simple case of constant competition matrix *K*_*ij*_ = *k*, competitive interactions are insensitive to the size of participating groups. In Appendix 1 of [Pichugin and Traulsen, 2020], we have shown that in this case, the equilibrium state **x**\* is proportional to the leading eigenvector of the projection matrix **A** - as in stationary regime of simpler model of growth without competition, compare Fig. 2A and B for an illustration.

**Figure 2:**
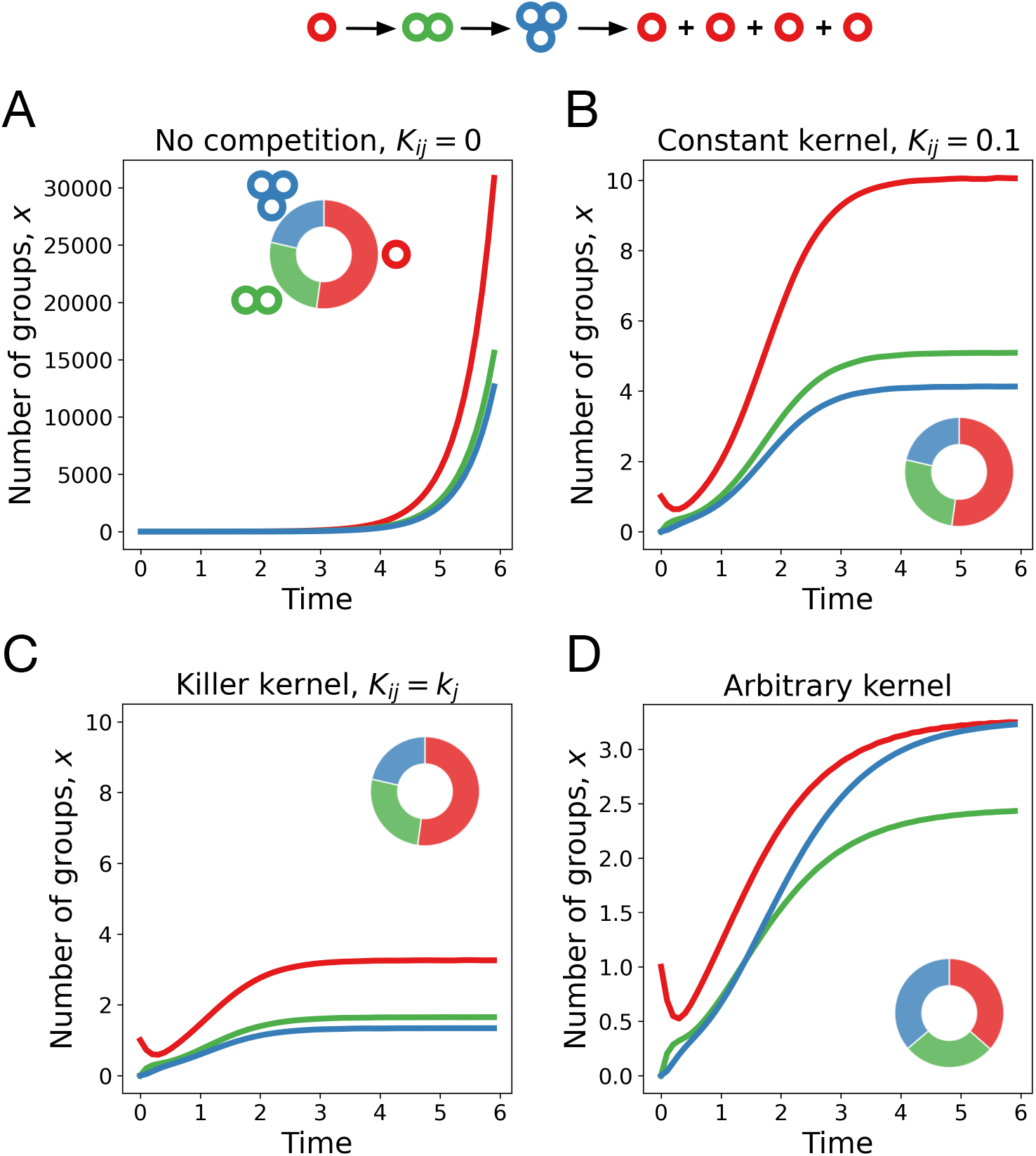
Example of the demographic dynamics of the life cycle 1+1+1+1 at different competition matrices. **A**. Without competition, the population grows exponentially, while the demographic distribution approaches a stationary composition (shown as a pie chart). Lines and wedges of different colours represent cell groups of different sizes: solitary cells (red), 2-celled groups (green), and 3-celled groups (blue). **B**. With a constant competition matrix, the population reaches a stationary state with the same demographic composition. **C**. Under a competition with a killer kernel, the demographic composition remains the same as in the linear model. **D**. For an arbitrary competition matrix, however, the demography at the equilibrium state can differ from the linear model. Figures are computed at *b*_*i*_ = (3, 2, 1), *d*_*i*_ = (0, 0, 0); for panel C, *K*_*ij*_ = *k*_*j*_, where *k*_*j*_ = (0.5, 0.1, 0.1); for panel D *K*_*ij*_ = *k*_*i*_, where *k*_*i*_ = (1, 0, 0).

The result above can be generalized: The stationary state **x**\* is proportional to the leading eigenvector of the projection matrix **A** not only for the constant competition but also for a broad class of competition matrices that we call a killer kernel,

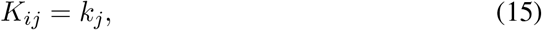

see Fig. 2C. There, the probability of a group to die in an encounter depends only on the size of the opponent group (*j*), hence the name. The proof that under an arbitrary killer kernel, the demographic composition in stationary state is the same as in the linear model is presented in Appendix A.1. However, for an arbitrary competition matrix, the composition of equilibrium state differs from the linear model, see Fig. 2D.

Similarly, we considered competition with a victim kernel, where the chance of a group to die depends only on the size of that group,

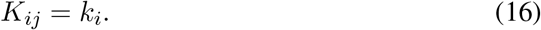

Here, we found another connection with the linear model. For competition with an arbitrary victim kernel, the total number of groups in population at equilibrium 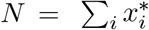 is equal to the growth rate of this population in the linear model of the same life cycle experiencing modified cell division rates 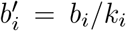 and group death rates 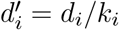, see Appendix A.2 for a proof.

### 3.2 Competition between multiple life cycles

A mixture of several sub-populations each executing a different life cycle, also reaches a stationary state. Some of the initially present life cycles may go extinct and do not contribute to the final population. Which life cycles remain in the population is a complex question. However, for special cases of interactions structure, this question can be addressed. For instance, in [Pichugin and Traulsen, 2020] we have shown that if the competition matrix is constant *K*_*ij*_ = *k*, then the outcome of evolution is identical to the linear model without group competition *K*_*ij*_ = 0 – only one life cycle survives, the one with the maximal population growth rate (e.g. leading eigenvalue of its projection matrix).

Here we found that the same result applies to any killer kernel (*K*_*ij*_ = *k*_*j*_). In this case, the composite competition matrix 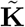 defined in Eq. (9) is a killer kernel as well. Since the population dynamics of both single and multiple life cycles are governed by equations with the same structure (Eqs. (2) and (7), respectively), the results of Appendix A.1 carry over to a composite population. Specifically, the stationary state of the composite population is proportional to the eigenvector of the composite projection matrix **Ã** corresponding to its leading eigenvalue. The composite projection matrix **Ã** defined in Eq. (8) is a block diagonal matrix, composed of projection matrices of the life cycles constituting the composite population. Thus, the leading eigenvalue of **Ã** is the largest among eigenvalues of all projection matrices comprising **Ã**. This rule is equivalent to the choice of the fastest growing life cycle in the linear model. Additionally, the corresponding eigenvector has non-zero components only at the positions in 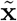 associated with the block having the maximal leading eigenvalue - i.e. only that life cycle survives to the stationary state. Thus, the outcome of life cycle evolution competing by a killer kernel (*K*_*ij*_ = *k*_*j*_) is exactly the same as in the linear model with unconstrained growth – the life cycle with the maximal growth rate outcompetes all others.

In the case of a victim kernel (*K*_*ij*_ = *k*_*i*_), also only a single life cycle survives in the long run, see the proof in Appendix B. There, each sub-population grows in numbers if the total population size is smaller than the equilibrium size (derived in Appendix A.2), and if the population size exceeds this value, the sub-population gradually dies out. Hence, selection favours the life cycle with the largest population size at the stationary state, because it can grow in a dense population, when all other life cycles die from intense competition.

### 3.3 Invasions from rare for a pair of life cycles

In the case of an arbitrary competition matrix **K**, the outcome of population dynamics does not always reduce to the dominance of a single life cycle as in the examples considered in the previous section. To investigate a scenario of an arbitrary competition matrix, we focus on a pair of life cycles (*κ*^(1)^, *κ*^(2)^) with the special initial conditions, where one of these life cycles is abundant, while the other one is rare. The life cycle *κ*^(1)^ can invade from rare into the abundant *κ*^(2)^ if the largest eigenvalue of the invasion matrix **A**^(1,2)^ is positive, see Eq. (12). Otherwise, the life cycle *κ*^(1)^ goes extinct.

Life cycle *κ*^(1)^ dominates life cycle *κ*^(2)^ if *κ*^(1)^ spreads from rare, but *κ*^(2)^ does not. This is equivalent to the leading eigenvalue of the invasion matrix **A**^(1,2)^ being positive, while the leading eigenvalue of **A**^(2,1)^ is negative. Then, independently of the initial conditions, only the life cycle *κ*^(1)^ survives in the long run, see Fig. 3A. The opposite signs results in the dominance of *κ*^(2)^ over *κ*^(1)^, see Fig. 3B.

**Figure 3:**
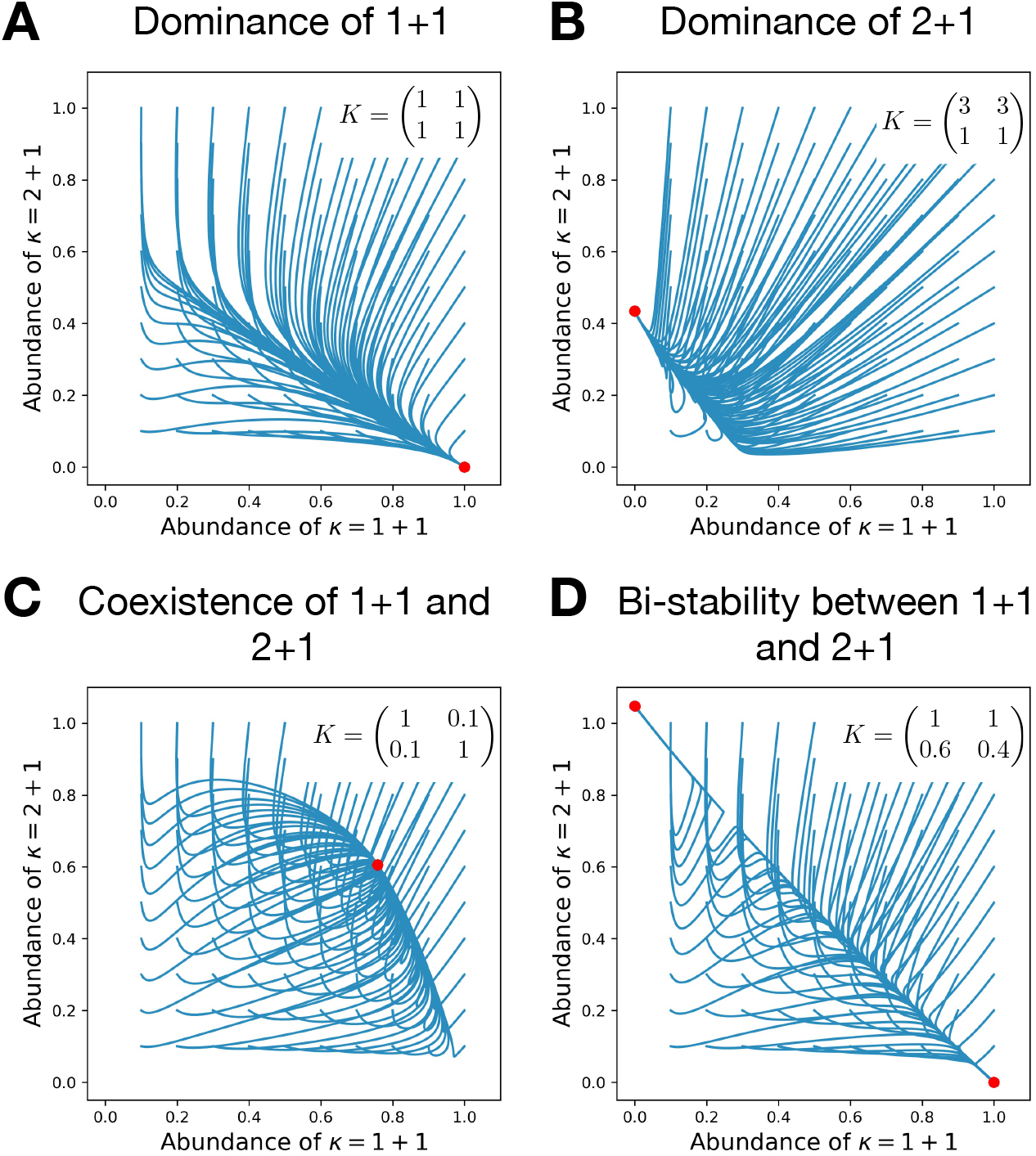
Competitive interactions can lead to the dominance of either life cycle, their coexistence, or bi-stability. Each panel shows the evolutionary trajectories of populations with various initial abundances of the life cycles 1+1 and 2+1 (blue lines) and all final states (red dots). Birth and death rates are *b*_*i*_ = (1, 0.5) and *d*_*i*_ = (0, 0), they favour the unicellular life cycle 1+1 in the absence of competition. Simulations performed on each panel only differ in the competition matrix **K**.

Beyond a dominance of one life cycle over another, it is possible that both life cycles are able to spread initially in the presence of the other. This happens when the leading eigenvalues of both invasion matrices **A**^(1,2)^ and **A**^(2,1)^ are positive. There, the result of interactions between life cycles is a coexistence of both - an outcome impossible in the model without competition, see Fig. 3C.

Finally, the leading eigenvalues of both invasion matrices could be negative – then neither of life cycles can invade into another. Then, the result is a bi-stability between life cycles, where the outcome of interactions depends on the initial conditions – another result impossible in the model without competition, see Fig. 3D.

The competition between groups plays a major role in determining which of four invasion patterns occurs. For instance, it is possible that a life cycle having an advantage in the raw growth rate (i.e. dominating in the linear model) is dominated as the result of competition, see Fig. 3B, where the life cycle 1+1 has a faster growth but 2+1 still dominates due to the advantage of multicellular groups in competition.

The leading eigenvalues of the invasion matrix can typically only be found numerically. First, the invasion matrix depends on the demographic composition of the resident population, which comes from a solution of a system of non-linear equations (10). Second, finding the eigenvalue requires solving a polynomial of degree equal to the size of the invasion matrix. However, one of the biologically most interesting scenarios – the evolution of multicellular life cycles in a population dominated by unicellular beings – deals with the most simple life cycle: 1+1. Therefore, some analytical results can be obtained here. An arbitrary multicellular life cycle *κ*^(*M*)^ can spread from rare in unicellular resident if this life cycle has a positive growth rate in a linear model with modified group death rates,

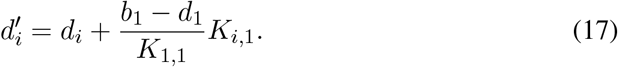

The successful invader drives the unicellular life cycle to extinction, if the unicellular life cycle cannot invade from rare, which occurs if

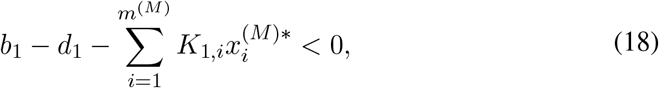

where *m*^(*M*)^ is the maximal group size in the life cycle *κ*^(*M*)^, and 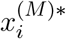 is the composition of the population when *κ*^(*M*)^ is abundant. Derivations of both conditions are presented in Appendix C

### 3.4 Invasions from rare for multiple life cycles

For multiple competing life cycles, the pattern of which life cycle can invade which can be very complex. Among this complexity, there is a special class of life cycles which cannot be invaded by any invader. These life cycles represent an evolutionary stable strategy (ESS) – because once such a life cycle dominates the system, it cannot be replaced by any other. Unlike in the linear model, an evolutionary stable strategy may not exist at all (then the stationary state is a coexistence of multiple life cycles) or more than one evolutionary stable strategy may be present (then different initial conditions may lead to different stationary states).

To illustrate the diversity of invasion patterns, we consider pairwise invasions in a triplet of life cycles *κ*^(1)^, *κ*^(2)^, *κ*^(3)^. The life cycle *κ*^(1)^ has 4 variants of stability against invasions: (i) either it is stable against invasions from both *κ*^(2)^ and *κ*^(3)^ (then *κ*^(1)^ is an evolutionary stable strategy), or (ii) stable against only *κ*^(2)^, or (iii) stable against only *κ*^(3)^, or (iv) both *κ*^(2)^ and *κ*^(3)^ can invade. The same four variants are possible for the two other life cycles. As a result, for the whole triplet, there are 4^3^ = 64 possible pairwise invasion patterns, which could feature 0, 1, 2, or 3 evolutionary stable strategies. Numerical simulations show that all 64 patterns can be expressed for the same triplet of life cycles given a right combination of cell birth, group death, and competition rates, see Fig. 4A.

**Figure 4:**
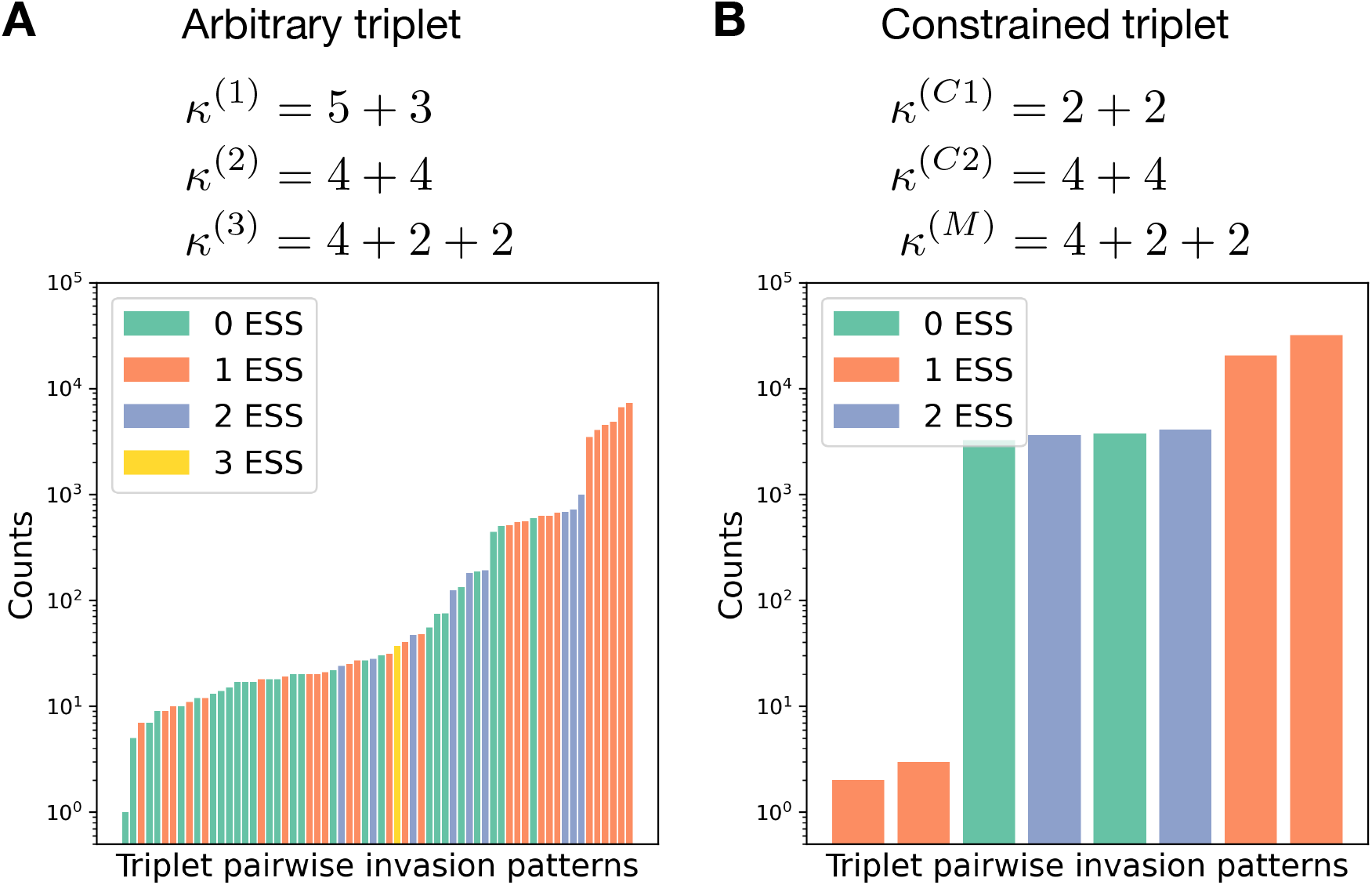
Constrained triplets demonstrate fewer patterns of pairwise invasion. **A**. For a combination of three life cycles, there are 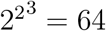 pairwise invasion patterns possible. For the triplet *κ*^(1)^ = 5 + 3, *κ*^(2)^ = 4 + 4, *κ*^(3)^ = 4 + 2 + 2, the rates of all processes (*b*_*i*_, *d*_*i*_, *K*_*i,j*_) were randomly sampled from an exponential distribution with unit rate parameter. Then the pairwise invasion pattern was identified. All 64 possible patterns were observed in these simulations. **B**. In a similar investigation for the triplet *κ*^(*C*1)^ = 2 + 2, *κ*^(*C*2)^ = 4 + 4, *κ*^(*M*)^ = 4 + 2 + 2, in which *κ*^(*C*1)^ and *κ*^(*C*2)^ constrain *κ*^(*M*)^, only eight patterns were found, see the main text for a discussion.

We generated a set of 40000 randomly drawn combinations of these rates from an exponential distribution with unit rate parameter and analysed the pairwise invasion patterns for each. The six most frequent patterns, comprising 77% of the generated data set feature a hierarchical dominance, where life cycles can be ordered in a way that higher order life cycle dominates (always invade) lower order life cycles. These six patterns are all possible hierarchical dominance triplets, as there are exactly six ways how three items can be placed in order. If we use the same analysis for a linear model, where life cycles compete for the growth rates, we will only observe hierarchical dominance as larger growth rate there results in domination. On the opposite side of the frequency spectrum, the two most rare patterns feature cyclic dominance, together comprising only 0.015% of dataset. There, in any pair of life cycles one dominates another but the whole triplet follows a “rock-paper-scissors” dynamics with no evolutionary stable strategies present (cf. [Park et al., 2020] studying why cyclic dominance is rare).

While an arbitrary triplet of life cycles may demonstrate up to 64 invasion patterns, some triplets, which we will call “constrained”, feature much smaller diversity of patterns. The triplet is constrained if the fragmentation rule of one (constrained) life cycle can be represented as a combination of fragmentation rules of two other (constraining) life cycles. The simplest example is the triplet *κ*^(*C*1)^ = 2 + 1, *κ*^(*C*2)^ = 1 + 1, *κ*^(*M*)^ = 1 + 1 + 1, where the fragmentation of three-celled group into three solitary cells (3 → 1 + 1 + 1) can be presented as a combination of the detachment of a singe cell (3 → 2 + 1) immediately followed by the dissolution of the two-cell group (2 → 1 + 1). A lot of constrained triplets can be constructed, e.g. in our illustrations we use the triplet *κ*^(*C*1)^ = 2 + 2, *κ*^(*C*2)^ = 4 + 4, *κ*^(*M*)^ = 4 + 2 + 2. Originally, constrained triplets emerged in the linear model [Pichugin et al., 2017], where the growth rate of the constrained life cycle *κ*^(*M*)^ was found to be always between growth rates of the constraining life cycles *κ*^(*C*1)^ and *κ*^(*C*2)^. It follows that in the model with competition (*K*_*ij*_ ≠ 0), each of two constraining life cycles (*κ*^(*C*1)^ and *κ*^(*C*2)^) must be either stable against two others, or unstable against both. The constrained life cycle *κ*^(*M*)^ in turn is always invaded by one of constraining life cycles and is stable against another, see Appendix D. Hence, the number of possible pairwise invasion patterns for such a triplet is limited to 2 · 2 · 2 = 8, see Fig. 4B. Among these eight patterns, two feature hierarchical dominance, where the life cycles can be ordered in a way that a higher order life cycle dominates the lower order life cycles (with the constrained life cycle *κ*^(*M*)^ always being in the middle of hierarchy), see Fig. 5A. In a larger dataset (66000 entries) with random birth, death, and competition rates, hierarchical dominance was observed in about 78% of entries. Two patterns feature bi-stability between constraining life cycles *κ*^(*C*1)^ and *κ*^(*C*2)^, see Fig. 5B. These patterns appear in about 11% of dataset. Two more patterns feature a coexistence between all three life cycles, see Fig. 5C. Note unlike to a pair of life cycles with unique coexistence equilibrium, the triplet features a range of stable coexistence states. The coexistence pattern is similarly frequent, observed in 11% of dataset. Finally, the least frequent two patterns are non-hierarchical dominance, where one constraining life cycle dominates another but in the abundance of the constrained life cycle, the invadeability is reversed, see Fig. 5D. They appear with three orders of magnitude lower frequency, smaller than 0.01% of cases.

**Figure 5:**
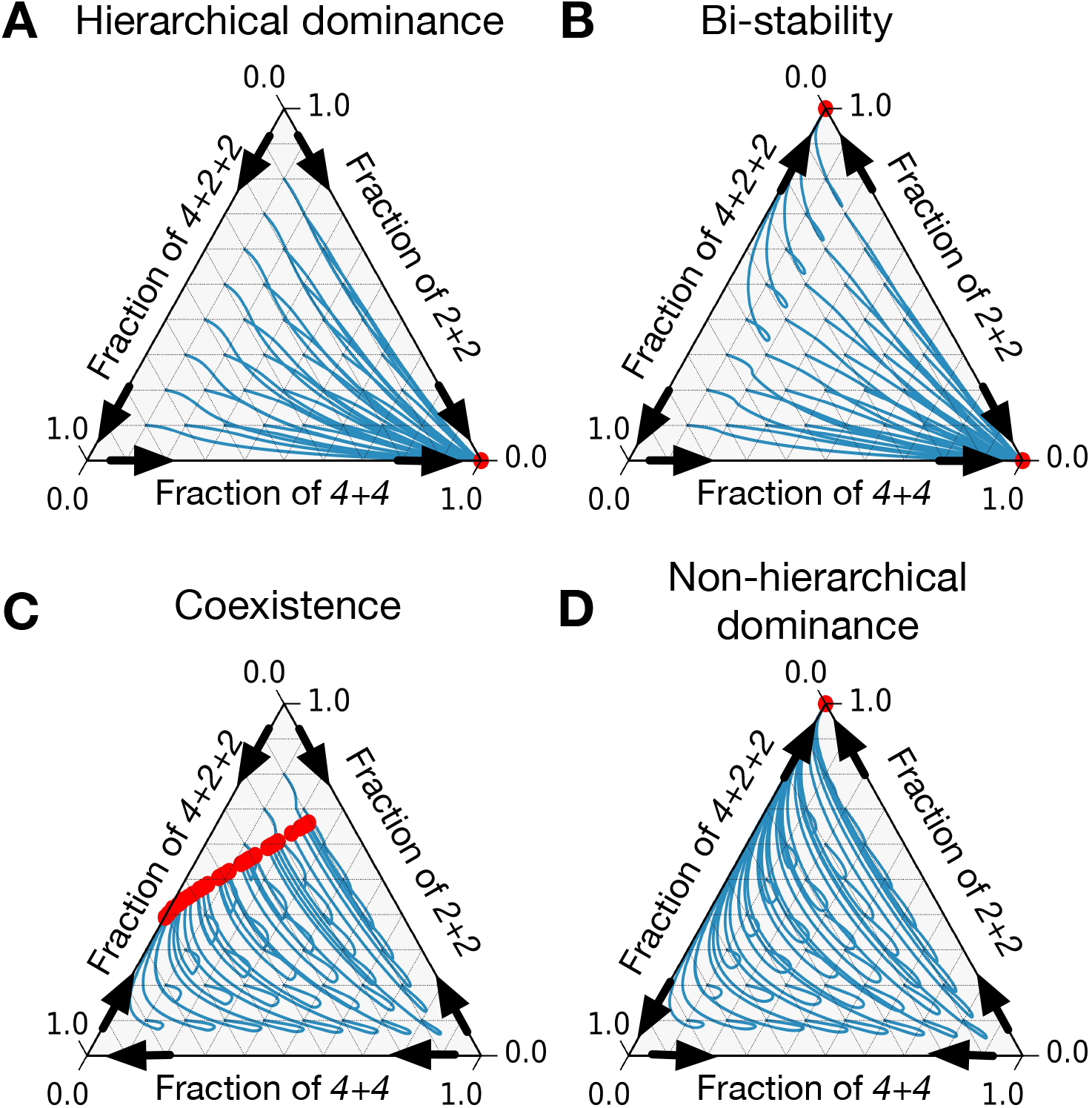
There are only eight patterns of pairwise invasion in a constrained triplet of life cycles. Four patterns are shown here and four more are symmetric to them. Blue lines are population dynamics trajectories with different initial composition of the population. Red points are final states. Black arrows show directions of invasion from rare. **A**. In a hierarchical dominance, one of constraining life cycles outcompetes the two others from any initial condition. **B**. In a bi-stability situation, each of two constraining life cycles is able to outcompete the third life cycle. Which of them will survive depends on the initial conditions. **C**. In a coexistence, all three life cycles survive to a stationary state. **D**. In a non-hierarchical dominance, one of constraining life cycles outcompetes another, however in the abundance of the constrained life cycle, the dominance is reversed. The parameters used to produce these figures are presented in appendix E.

The analysis of constrained triplets above shed light on the evolutionary dynamics when more than three life cycles are involved. For instance, in the linear model, the existence of constrained triplets immediately means that evolutionarily optimal life cycles must be a fragmentation of a group into two pieces. In a constrained triplet, both constraining life cycles (*κ*^(*C*1)^ and *κ*^(*C*2)^) have smaller number of offspring then the constrained life cycle (*κ*^(*M*)^). Hence, for any life cycle and for any birth (*b*_*i*_) and death (*d*_*i*_) rates, there is a life cycle with smaller number of offspring and a larger population growth rate, unless the life cycle is binary (which cannot be constrained). Similarly, in our present eco-evolutionary model, the constrained life cycle (*κ*^(*M*)^) can always be invaded by exactly one constraining life cycles (*κ*^(*C*1)^ and *κ*^(*C*2)^). Thus, under arbitrary combination of birth (*b*_*i*_), death (*d*_*i*_), and competition (*K*_*i,j*_) rates, any life cycle can be invaded by at least one other life cycle, in which groups fragment into a smaller number of pieces, see Appendix D for the proof. As a result only binary fragmentation life cycles can be evolutionary stable strategies (if all life cycles are initially present / can eventually emerge in the population).

## 4 Discussion

In this work, we have added an ecological dimension to the problem of life cycle evolution. We asked the question what clonal life cycles can evolve when groups compete with each other – for a limited resource or space.

Here, we considered only costless fragmentation. If fragmentation is costly (e.g. imposes some risk of cell loss), binary fragmentation is no longer special, as multiple fragmentation modes are capable to win the growth rate competition [Pichugin and Traulsen, 2020]. Nevertheless, some “forbidden” life cycles remain constrained. These are the life cycles containing two different subsets of offspring with the same combined size. For instance *κ*^(*M*)^ = 2 + 1 + 1 is forbidden as it has offspring subsets 2 and 1 + 1 with the same combined size. In [Pichugin and Traulsen, 2020], we have shown that the growth rate of this life cycle is constrained between the growth rates of life cycles *κ*^(*C*1)^ = 2 + 2 and *κ*^(*C*2)^ = 1 + 1 + 1 + 1, which use the same subsets twice. All our findings related to the constrained triplets hold true for these triplets as well. The results demonstrated in the present work for multiple fragmentation directly transfer to forbidden life cycles if fragmentation is costly.

In the broad context of the eco-evolutionary dynamics, our dynamical equations (2) and (7) bear a similarity with generalized Lotka-Volterra equations: both contain a linear growth term and a non-linear competition term (typically of the second order) balancing out the linear growth. However, our equations are not equivalent to the generalized Lotka-Volterra (GLV). In the GLV, individuals corresponding to different elements of the population vector reproduce independently, i.e. in our terms, the projection matrix **A** is diagonal. In our model, however, an individual group changes its state in the course of life cycle and the projection matrix **A** is not diagonal. Yet, it is possible to introduce a linear transformation **x** → **Cy**, where **C** is a matrix, which will make the linear term in our model diagonal. However, in this case, the interaction term will loose the GLV form of the modification of the growth rate (−*x*_*i*_*Σ*_*j*_ *K*_*ij*_*x*_*j*_, see Eq. (1)) and will become a general second-order term instead (−Σ_*jk*_ *K*_*ij*_*x*_*j*_*x*_*k*_). Given that our system is not a generalized Lotka-Volterra in disguise, it surprising how much of the analysis presented here has been performed using the approaches designed to treat GLV systems.

In the context of the theory of the evolution of multicellularity (e.g. [Roze et al., 2001, Tarnita et al., 2013, Pichugin et al., 2017]), our work extends past work by introducing a competition between cell groups, which leads to a more complicated non-linear model. Here, neither previously employed analysis can be applied (there is no “fitness” measure to optimize), nor the results can be directly compared (there is no guarantee of the existence of the single optimal life cycle). Hence, from the broader perspective of life cycle evolution theory, two questions arise:

1. What aspects of life cycles evolution were correctly captured by models neglecting group competition?
2. What aspects of life cycles evolution are impacted by taking into account competition between groups?

### Similarities between models with and without group competition

Comparing our results with growth competition models [Pichugin et al., 2017, Pichugin and Traulsen, 2020], we found that they are capable to capture short-term dynamics of the evolution. While the population composition changes a little, the present model is effectively a growth competition with modified death rates, see Appendix D. Thus, the approach of fitness optimization, used in simpler models, is adequate to the description of evolutionary stable strategies, for which population composition is resilient to perturbations. It follows that in the present model, the evolutionary stable strategy must be some sort of binary fragmentation, where a parent group splits into exactly two offspring – the same class of life cycles restricts the optimal strategies in the growth competition model [Pichugin et al., 2017].

We also identified two classes of competition matrices (killer kernel and victim kernel), in which the evolution outcome can be inferred from the model without group interactions. Under the killer kernel, where the death probability only depends on the size of the opponent group, the eco-evolutionary dynamics favours life cycles with the largest population growth rate. Under the victim kernel, where the death probability only depends on the size of the group dying, the eco-evolutionary dynamics favours life cycles with the largest carrying capacity.

Simulations of pairwise invasion patterns revealed additional similarities between two models. The model without group competition has a measure of life cycle fitness – the population growth rate, and all life cycles can be ranked by its value. Therefore, the pairwise invasion pattern is always a hierarchical dominance: more fit life cycles always invade less fit ones. Numerical simulations of our model with group competition have shown that the majority of randomly sampled competition matrices result in the same invasion pattern of the hierarchical dominance, see Fig. 4. However, it remains unknown if any measure of life cycle fitness can be defined for any of these cases.

### Differences between models with and without group competition

Our current model taking into account the competition between groups demonstrates a different and more rich long-term dynamics than simpler models neglecting such a competition. Not only group competition can lead to the success of the life cycle with a slower growth rate (Fig. 3B) but also allow for more complex outcomes of evolution. First, a stable coexistence of multiple life cycles is possible, see Fig. 3C. As a consequence, multiple fragmentation life cycles can survive as a part of mixed population, see Fig.5C. A similar result was observed in a model with dynamic environments [Pichugin et al., 2019], where a multiple fragmentation life cycle was observed as a component of an evolutionarily optimal mixed life cycle. Second, the outcome of evolution may depend on the initial state of population, see Fig. 3D. Hence, there might be one, none, or several evolutionary stable strategies.

## Conclusion

Summarizing the comparison above, models neglecting group competition capture well the short-term dynamics and features of evolutionary stable strategies. However, they fall short in the description of long-term dynamics and outcomes on the population-wide scale.

## A Dynamics of a single life cycle

### A.1 Population dynamics under the killer kernel

In this section, we show that the demographic distribution of populations in a stationary regime is identical in the linear model (*K*_*i,j*_ = 0) and under the killer kernel (*K*_*i,j*_ = *k*_*j*_).

In the linear model, the stationary state is an exponentially growing population [Pichugin et al., 2017]

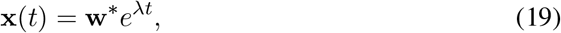

where *λ* is the leading eigenvalue of the matrix **A** and **w**\* is the corresponding eigen-vector,

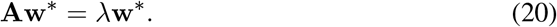

Under the killer kernel, death rates due to competition are the same for groups of all sizes. Hence, following Eq. (2), the population dynamics of a life cycle under a killer kernel is

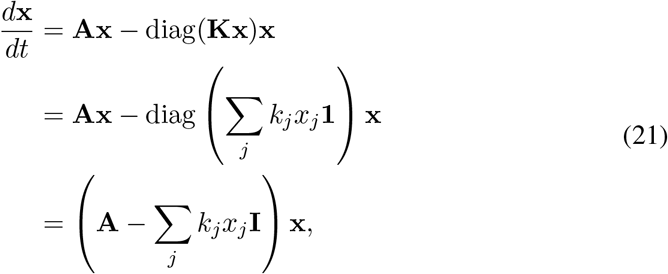

where **1** is the vector of ones, and **I** is the identity matrix (diag(**1**) = **I**).

In can be shown by a direct calculation, that the vector

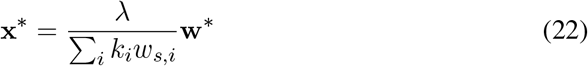

is the stationary state of the dynamics (21):

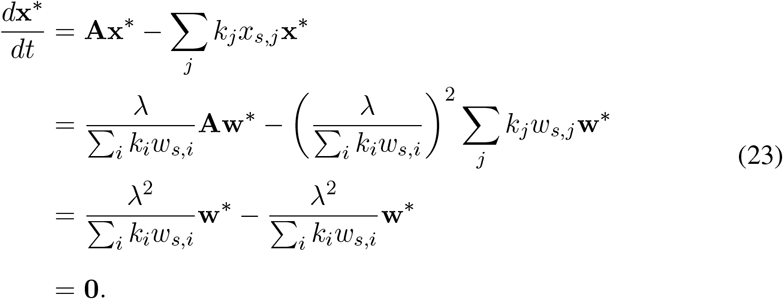

Since the stationary state **x**\* under the killer kernel is proportional to the vector describing the stationary distribution **w**\* in the linear model, the population composition in both scenarios are the same.

### A.2 Population dynamics under the victim kernel

In this section, we show that the task of finding the equilibrium population size 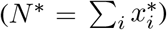 under the victim kernel (*K*_*i,j*_ = *k*_*i*_) is mathematically equivalent to the task of finding the population growth rate in the linear model (*K*_*i,j*_ = 0) with modified cell birth rates (*b*_*i*_ → *b*_*i*_*/k*_*i*_) and group death rates (*d*_*i*_ → *d*_*i*_*/k*_*i*_).

In the linear model, the population growth rate is found as the leading eigenvalue of the projection matrix determined by [Pichugin et al., 2017]

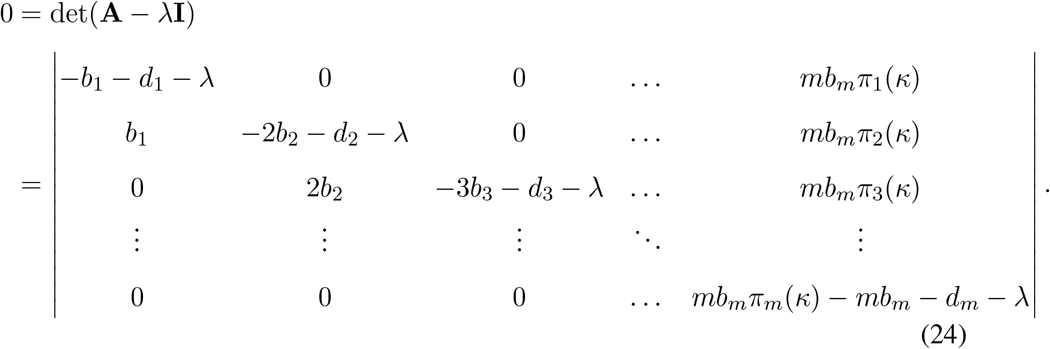

Under the victim kernel, the death rate due to the competition depends only on the size of the outcompeted group. Hence, following Eq. (2), the stationary state of a life cycle under a victim kernel satisfies

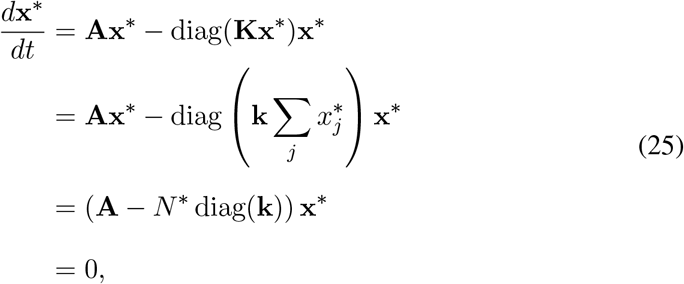

where **k** is a vector constructed from any column of the competition matrix **K** (they are all identical), and 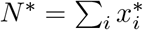 is the equilibrium population size.

The last equality in Eq. (25) implies that by Fredholm alternative, one out of two xconditions is satisfied [Hoffman, 1971]:

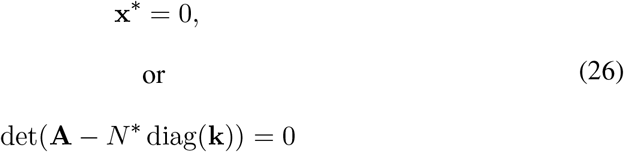

Limiting ourselves to scenarios where the stationary state **x**_0_ is not an empty population, we can conclude that the population at the equilibrium satisfies

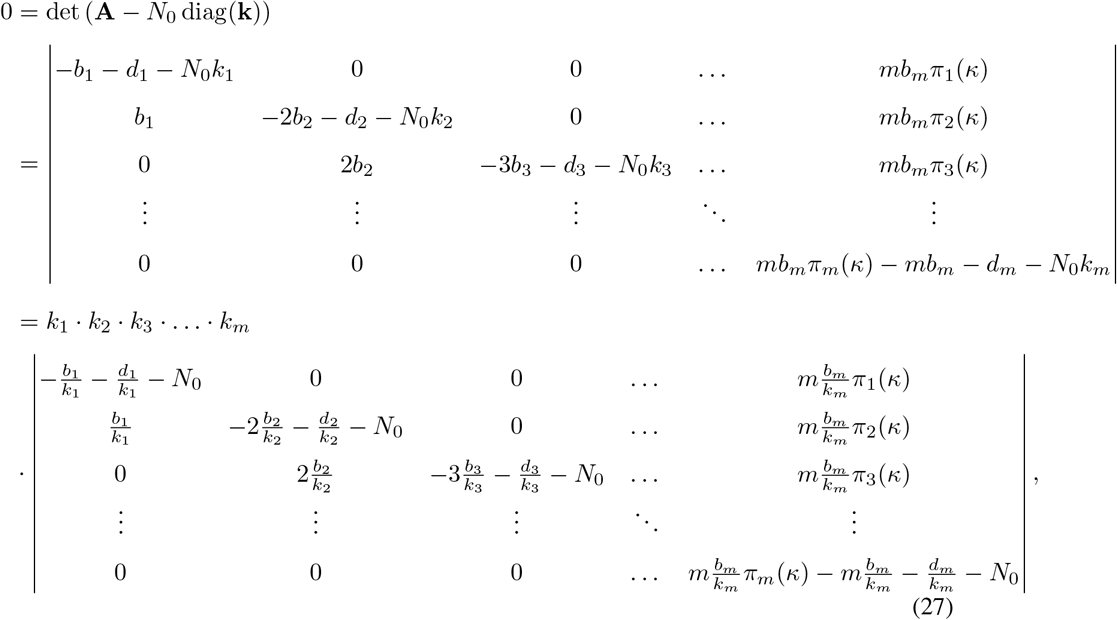

where in the last step, we divided the *i*-th column by *k*_*i*_ for all *i*. Comparing the determinants in Eqs. (24) and (27), we find that they are identical after the substitution

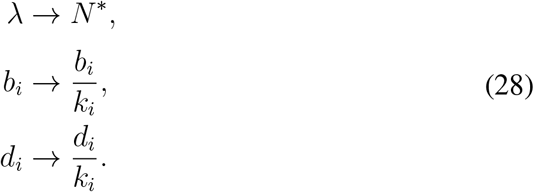

Thus, the equilibrium population size (*N*\*) under a victim kernel can be found as a population growth rate in the linear model with modified cell birth and group death rates.

## B Competition under the victim kernel

In this section, we show that in a composite population, in which multiple (*r*) life cycles compete through a victim kernel (*K*_*i,j*_ = *k*_*i*_), only one life cycle survives to the stationary state, and this is the same life cycle that has the maximal carrying capacity if grown in isolation, and the same life cycle that would be evolutionarily optimal in the linear model (*K*_*i,j*_ = 0) with modified cell division (*b*_*i*_ → *b*_*i*_*/k*_*i*_) and group death (*d*_*i*_ → *d*_*i*_*/k*_*i*_) rates.

If the competition matrix **K** constitutes a victim kernel, then the composite competition matrix 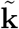 defined as in Eq. (9) is a victim kernel as well. Hence, the results from Appendix A.2 hold also for a composite population. In particular, the population size at the stationary state 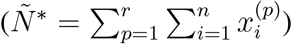 is determined by

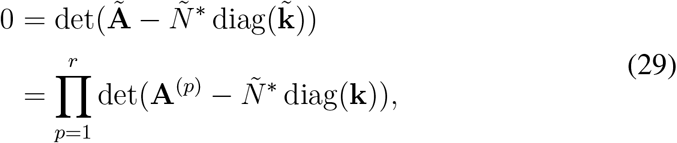

where 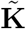 is the vector made from a single column of the composite competition matrix 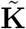 (it is a concatenation of *r* vectors **k**). In the second step of Eq. (29), we used that the determinant of a block-diagonal matrix is the product of determinants of its blocks. Eq. (29) is satisfied if one of the multipliers is equal to zero. The *p*-th multiplier becomes zero, when the composite population reaches the size equal to the carrying capacity of the *p*-th life cycle, cf. Eq. (26). At that moment, the *p*-th life cycle can neither grow or decay, the life cycles with lower carrying capacity decay as they cannot keep up with competition caused by overcrowding, and only the life cycles with carrying capacity larger than the population size can grow in numbers. Eq. (29) has *r* possible solutions with respect to *Ñ*\* - one for each competing life cycle. However, only one solution can represent a stationary population, where no life cycle can grow in numbers – the one with the maximal population size. There, one life cycle is stationary, while all others decrease in numbers due to overcrowding. Thus, the outcome of life cycle competition under a victim kernel is the survival of a single life cycle, which has the maximal equilibrium population size among all competitors. According to Appendix A.2, these population sizes are equal to the growth rates of the corresponding life cycles in a linear model with the modified cell division and group death rates. Consequently, the maximal population size corresponds to the fastest growing life cycle in that modified linear model. The population corresponding to the highest eigenvalue takes over and will dominate the system. This means that the life cycle having the largest population size in isolation dominates all other life cycles in the competition through a victim kernel.

## C Invasion into unicellular resident and invasion of the unicellular invader

If the resident is unicellular (*κ*^(*R*)^ = 1+1), its steady state is given by the solution of

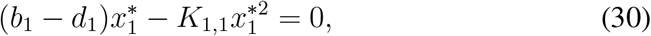

equal to

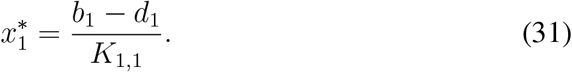

Then, for an arbitrary invader multicellular life cycle *κ*^(*M*)^, the invasion matrix is given by

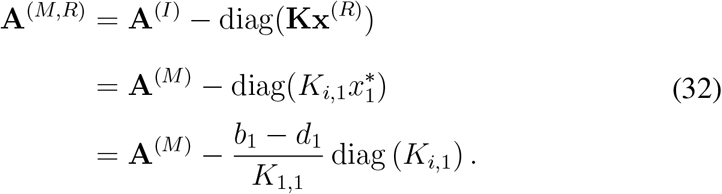

This is equivalent to the linear growth of the invader life cycle in an environment with modified death rates

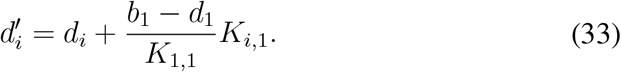

If the invader is unicellular (*κ*^(*I*)^ = 1 + 1), then the invasion matrix is effectively a 1 × 1 matrix, since the invader contains only isolated cells. Even if **A**^(*I,M*)^ formally has a larger size, it is a block matrix, with the element 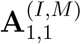 being a block, with a value that is equal to the growth rate of the unicellular life cycle. This value is

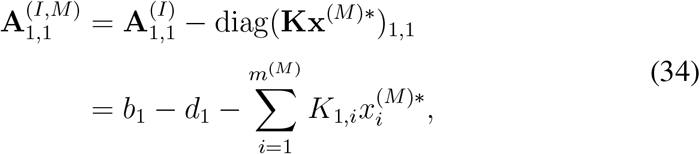

where 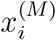 is the number of groups of size *i* in the resident population and *m*^(*M*)^ is the maximal group size of the resident life cycle. The unicellular invader cannot spread in a resident population, when 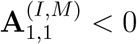.

## D Invasion of life cycles and restrictions on ESS

In this section, we show that any resident life cycle with fragmentation into multiple parts can always be invaded by at least one life cycle with a smaller number of offspring.

Consider a resident life cycle *κ*^(*R*)^, in which more than two offspring groups are produced in the result of fragmentation. The initial dynamics of any life cycle *κ*^(*I*)^ invading from rare, can be described by a linear model with death rates modified as

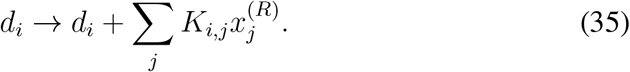

The invasion is successful if the leading eigenvalue of the corresponding projection matrix **A**^(*I,R*)^, defined as in Eq. (11), is positive.

In the analysis of the linear model [Pichugin et al., 2017], it was shown that if the fragmentation preserves the number of cells (no cell loss), then for any multiple fragmentation life cycle, there exist two constraining life cycles with a smaller number of offspring. For any combination of cell division rates (*b*_*i*_) and group death rates (*d*_*i*_), one of the constraining life cycles has a larger growth rate than the focal multiple fragmentation life cycle, while another has a smaller growth rate.

Now consider the invasion from rare in our present model with competition. The initial invasion rate, computed to the leading eigenvalue of the matrix **A**^(*I,R*)^ is equal to the growth rate of the invader life cycle in an environment modified according to Eq. (35). Therefore, for any resident population, the invasion rate of the constrained life cycle is always in between the invasion rates of the constraining life cycles. If the resident population is formed by the constrained life cycle itself, its self-invasion rate is zero. Hence, one of the constraining life cycles has a larger invasion rate (i.e. it is positive), while another has a smaller invasion rate (negative). As a result, the constrained life cycle can always be invaded by exactly one of its constraining life cycles. No constrained life cycle can be an evolutionary stable strategy. Since any life cycle with more than two offspring is constrained, only binary fragmentation can be an evolutionary stable strategy.

To conclude this appendix, we consider the resident population formed by a constraining life cycle. If the constrained life cycle has positive invasion rate, then another constraining life cycle must have a positive invasion rate as well. Alternatively, the constrained life cycle has negative invasion rate, then another constraining life cycle also has negative invasion rate. Thus, a constraining resident is either resistant to invasion from both other life cycles in a triplet, or can be successfully invaded by both of them.

## E Parameters of calculation used in figures

In Fig. 2, we used *b* = (3, 2, 1) and *d* = (0, 0, 0). In all cases, the population was initialized identically *x*|_*t*=0_ = (1, 0, 0). The dynamics shown at various panels differ by the competition matrix used:

- Panel A

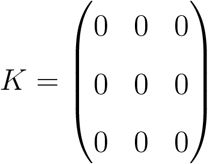
- Panel B

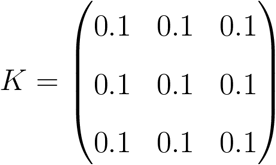
- Panel C

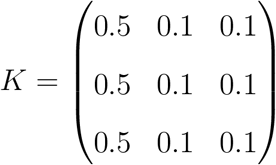
- Panel D

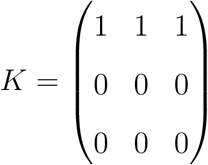

In Fig. 3, we considered the life cycles 1+1 and 2+1. We used *b* = (1, 0.5) and *d* = (0, 0). For each plotted trajectory, the populations were initialized with **x**_1+1_ = (*s*_1_), **x**_2+1_ = (*s*_2_, 0), where *s*_1_, *s*_2_ ∈ {0.1, 0.2, 0.3, …, 1.0}. The dynamics shown in the four panels differ by the competition matrix used:

- Panel A

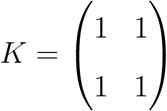
- Panel B

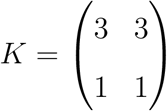
- Panel C

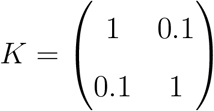
- Panel D

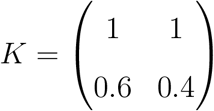

In Fig. 5, we considered the life cycles *κ*^(*C*1)^ = 2 + 2, *κ*^(*C*2)^ = 4 + 4, *κ*^(*M*)^ = 4 + 2 + 2. Panels differ in birth, death, and competition rates. Trajectories in each panel have different initial states. For each initial state, the composite population contains all three life cycles with different fractions *s*^(*C*1)^, *s*^(*C*2)^, *s*^(*M*)^, such that *s*^(*C*1)^ + *s*^(*C*2)^ + *s*^(*M*)^ = 1. The initial group sizes distribution is proportional to the equilibrium population of that life cycle alone:

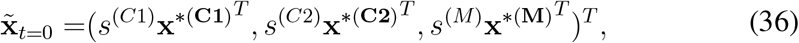

where the vectors 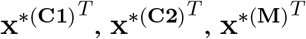 are equilibrium population states of corresponding life cycles grown in isolation, computed according to Eq. (14).

- In panel A (hierarchic dominance), we used *b*_*i*_ = 1.0, *d*_*i*_ = 0 and the competition matrix *K*_*ij*_ = 0.1.

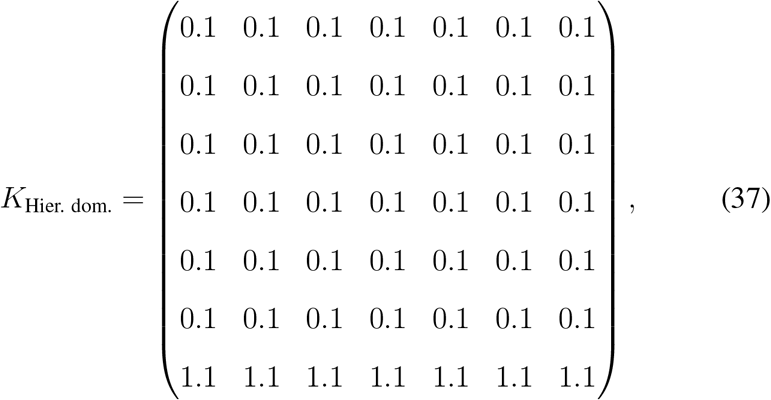

or equivalently

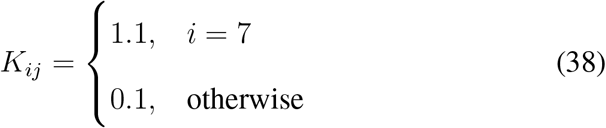
- In panel B (bi-stability), we used *b*_*i*_ = 1 and *d*_*i*_ = 0, while the competition matrix was

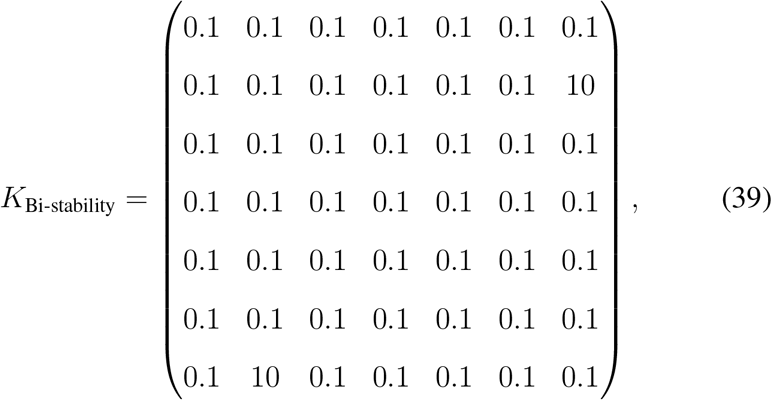

or equivalently

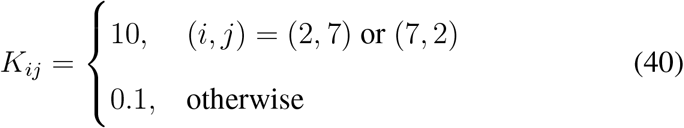
- In panel C (coexistence), we used *b*_*i*_ = 1 and *d*_*i*_ = 0, while the competition matrix was

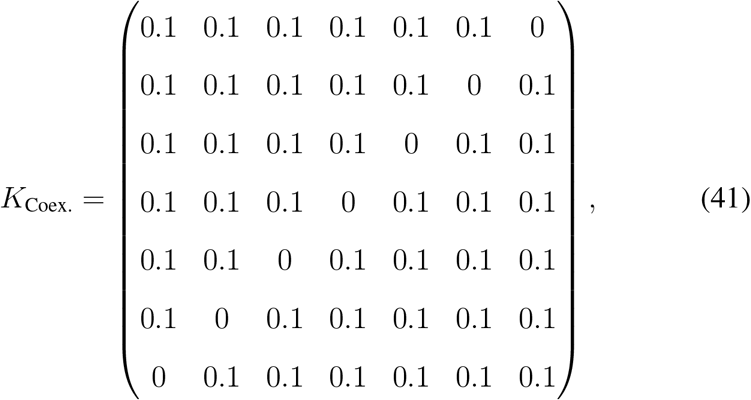

or equivalently

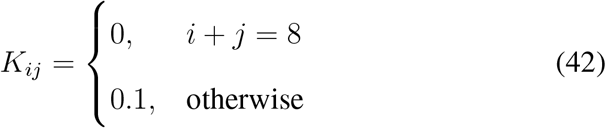
- In panel D (Non-hierarchical dominance), we used *d*_*i*_ = 0,

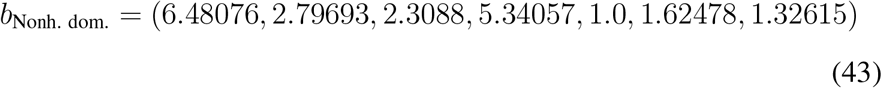

and

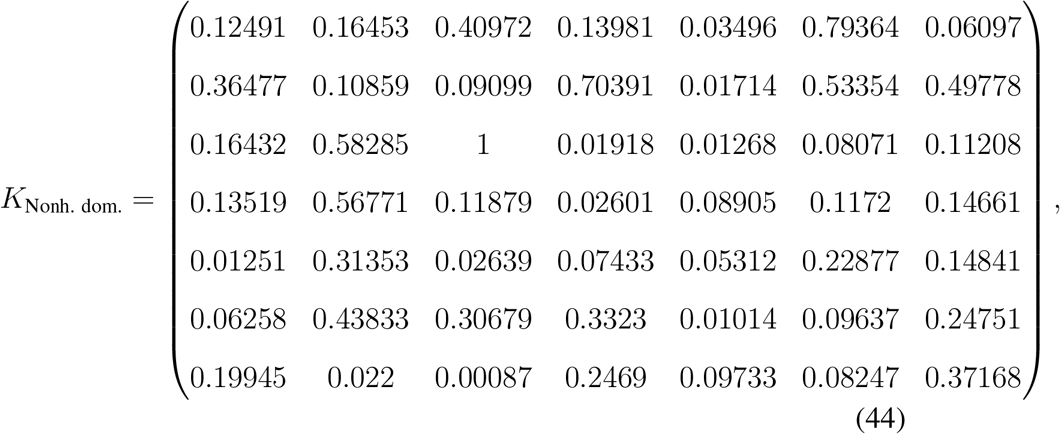

